# An atlas of robust microbiome associations with phenotypic traits based on large-scale cohorts from two continents

**DOI:** 10.1101/2020.05.28.122325

**Authors:** Daphna Rothschild, Sigal Leviatan, Ariel Hanemann, Yossi Cohen, Omer Weissbrod, Eran Segal

## Abstract

Numerous human conditions are associated with the microbiome, yet studies are inconsistent as to the magnitude of the associations and the bacteria involved, likely reflecting insufficiently employed sample sizes. Here, we collected diverse phenotypes and gut microbiota from 34,057 individuals from Israel and the U.S.. Analyzing these data using a much-expanded microbial genomes set, we derive an atlas of robust and numerous unreported associations between bacteria and physiological human traits, which we show to replicate in cohorts from both continents. Using machine learning models trained on microbiome data, we predict human traits with high accuracy across continents. Subsampling our cohort to smaller cohort sizes yielded highly variable models and thus sensitivity to the selected cohort, underscoring the utility of large cohorts and possibly explaining the source of discrepancies across studies. Finally, many of our prediction models saturate at these numbers of individuals, suggesting that similar analyses on larger cohorts may not further improve these predictions.

## Introduction

The human gut microbiota is linked to metabolic disorders such as diabetes and obesity but these links are based on relatively small cohorts of several dozens or hundreds of individuals ^1–8^. Although these studies reported many statistically significant associations, many of these effects are either moderate or do not replicate in other works ^9,10^. One such example is alpha diversity, for which there are contradicting reports regarding its association with different phenotypes. While microbiome diversity is mostly regarded as a positive indicator of health ^8,11–14^, other studies found that increased diversity is associated with microbiome instability ^15,16^. Diversity was also shown to increase with age ^17,18^, but this association was not conclusive in other cohorts ^19^. These discrepancies call for studying these questions across larger cohorts from diverse backgrounds as was done for several studies comparing the microbiome in different populations ^20–22^. Indeed, in the field of genetics, large cohorts are required since many traits are known to be polygenic and to be affected by small effects from many variants ^23,24^. Similarly, in the microbiome we expect that individual bacterial species may have a low abundance or mild associations with human phenotypes, necessitating large sample sizes. In addition, many bacterial species are present in only a relatively small fraction of the population such that the association between their abundance and traits can only be studied in large cohorts that have enough individuals that harbor them.

Apart from cohort size, there are other challenges in finding robust signals from the microbiome. One such challenge stems from the large number of genes that are shared between different bacteria through mechanisms such as horizontal gene transfer ^25,26^. Such sharing causes many short metagenomic sequencing reads to map non-uniquely to multiple bacteria, making it difficult to estimate bacterial relative abundance. Several methods were devised to address this issue, e.g., by mapping to genes that appear in a single copy and are unique to a single species ^27^. However, these methods need to be applied anew every time that we wish to use a different reference genome set, as we wished to do here given the much expanded reference of bacterial species groups (SGBs) published in 2019 which added 3,796 new SGBs to the human microbiome catalog ^28^. Another challenge stems from only being able to estimate microbial relative abundances, not absolute, which can lead to false bacteria-phenotype associations. This concern can partially be addressed with rarefaction samples and using *reference-frames*^*29*^, with best practice being the ability to replicate results on independent datasets.

To address the above issues and with the aim of deriving robust microbiome associations, we used metagenomic sequencing to profile the gut microbiome of 34,057 individuals from both Israel and the U.S., for which we also obtained a rich set of self reported phenotypes. We devised a novel algorithm for assessing bacterial relative abundances based on unique genetic elements, and applied it to the recent and much expanded SGB dataset of Pasolli et al. ^28^. Using the relative abundances on this expanded genome set and much larger cohort, we identified numerous associations between microbiome diversity and several human traits. We were also able to develop models that predict these traits using only microbiome data with high accuracy, as in the case of age (R^2^=0.31). Notably, these associations replicate across continents, and models derived from the Israeli cohort generalize well to the U.S. cohort, so they are not specific to a certain environment.

By subsampling our cohort to typical cohort sizes used in other studies, we show that associations and predictions derived from smaller cohorts are highly variable and thus sensitive to the selected cohort, underscoring the need for larger cohorts in the microbiome field.

## Results

### Metagenome samples for 34,057 participants from two continents

We obtained gut metagenomic profiles from 30,083 and 3,974 individuals from Israel and the U.S., respectively, who submitted their sample to a consumer microbiome company and signed an appropriate consent form. Participants also answered questionnaires and provided self-reported phenotypic data and blood tests (**Table 1**).

**Table 1.** Values are mean ± one std. Number of participants for which we have the phenotype is shown in parenthesis.

|  | **Israel train** | **Israel test** | **U.S. test** |
| --- | --- | --- | --- |
| **Age (years)** | 54.80±13.67<br>(n=27,075) | 54.62±13.50<br>(n=3,008) | 52.35±13.13<br>(n=3,974) |
| **Albumin (g/dl)** | 4.22±0.80<br>(n=14,098) | 4.19±0.79<br>(n=1,564) | 4.18±0.91<br>(n=2,018) |
| **Alkaline phosphatase (U/L)** | 79.75±130.61<br>(n=15,245) | 86.23±207.73<br>(n=1,720) | 68.70±113.53<br>(n=2,034) |
| **BMI (kg/cm<sup>2</sup>)** | 28.23±4.99<br>(n=27,018) | 28.36±5.11<br>(n=3,003) | 27.07±5.68<br>(n=3,951) |
| **Bowel movement (\*)** | 2.86±0.68<br>(n=26,953) | 2.86±0.68<br>(n=2,997) | 2.88±0.64<br>(n=3,944) |
| **Currently smokes Y:N** | 2,397:24,575 | 273:2,728 | 72:3,869 |
| **Ever smoked Y:N** | 10,606:16,366 | 1,172:1,829 | 1,017:2,924 |
| **Gender M:F** | 11,019:16,056 | 1,129:1,779 | 1,678:2,296 |
| **Glucose (mg/dl)** | 103.40±24.80<br>(n=21,613) | 103.62±25.24<br>(n=2,405) | 97.74±22.60<br>(n=2,271) |
| **HbA1C% - Hemoglobin A1C (%)** | 5.81±0.85<br>(n=19,457) | 5.80±0.84<br>(n=2,181) | 5.52±0.73<br>(n=2,292) |
| **HDL cholesterol (mg/dl)** | 53.15±17.72<br>(n=22,391) | 53.00±16.93<br>(n=2,507) | 61.09±20.93<br>(n=2,546) |
| **Height (cm)** | 168.68±9.12<br>(n=27,075) | 168.86±9.28<br>(n=3,006) | 170.36±10.30<br>(n=3,971) |
| **INR** | 1.13±1.22 | 1.17±1.41 | 1.42±1.91 |
|  | (n=7,034) | (n=809) | (n=488) |
| **Protein (g/dl)** | 7.13±1.21<br>(n=13,664) | 7.08±1.37<br>(n=1,520) | 6.80±1.44<br>(n=2,008) |
| **SGOT - serum**<br>**glutamic oxaloacetic**<br>**transaminase (IU/L)** | 25.87±62.92<br>(n=19,648) | 29.04±112.83<br>(n=2,182) | 24.80±42.76<br>(n=2,098) |
| **Total cholesterol**<br>**(mg/dl)** | 185.74±44.02<br>(n=21,965) | 185.45±43.25<br>(n=2,463) | 190.19±50.12<br>(n=2,532) |
| **Triglycerides (mg/dl)** | 131.62±78.27<br>(n=22,149) | 134.11±81.97<br>(n=2,484) | 107.12±70.94<br>(n=2,432) |
| **TSH - Thyroid**<br>**stimulating hormone**<br>**(microU/ml)** | 2.75±6.16<br>(n=14,958) | 2.58±4.20<br>(n=1,662) | 3.48±10.82<br>(n=2,016) |
| **Type 2 diabetes Y:N** | 4,369:22,746 | 479:2,529 | 294:3,680 |

(\*) Bowel movement is a categorical variable which takes the following values:
- 0 - Less Than Once A Week - 1 - One or Two Times Per Week - 2 - Three or Four Times Per Week - 3 - One or Two Times Per Day - 4 - Three to Five Times Per Day - 5 - More Than 5 Times Per Day

We randomly selected 90% of the samples from the Israeli cohort (n=27,075 samples) to be our discovery cohort on which we trained predictive models using cross-validation and set aside as independent test sets the remaining 10% of the Israeli cohort (“test1”, n=3,008 samples) and the entire U.S. cohort (“test2”, n=3,974 samples) (**Figure 1a-e, Table 1**). These test sets were only used once to evaluate the performance of the models developed on the discovery cohort.

**Figure 1:**
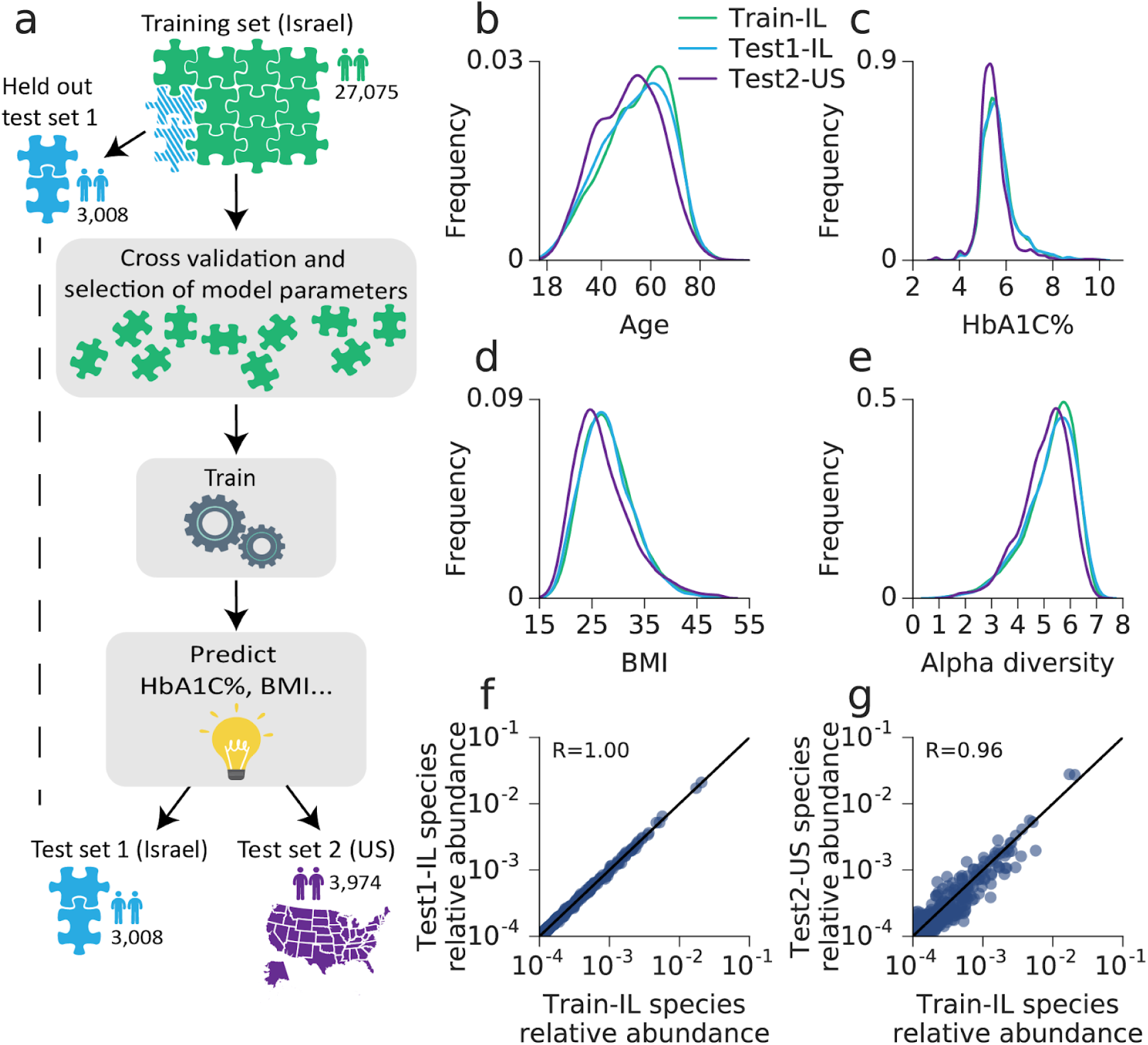
Cohort description and model prediction scheme. (a) Illustration of cohorts and machine learning process. A training set of 27,075 individuals was randomly selected out of 3,083 Israeli individuals and was used for model parameter selection using 10 fold cross validation and microbiome, age and gender features. For each phenotype the selected model was trained on the 27,075 training samples and then tested on both the held out 3,008 samples of the Israeli population and a separate U.S. test cohort of 3,974 individuals. (b) Distribution of age in the 3 cohorts, training test1 and test2. (c) - (e) Same for HbA1C%, BMI and alpha diversity. (f) A scatter plot comparing the mean log relative abundance of each species, in the Israeli training cohort vs. the Israeli test1 cohort. R value represents Pearson correlation. (g) Same as (f) in the Israeli training cohort vs. the US test2 cohort. R value represents Pearson correlation.

To compute bacterial relative abundance, we used the representatives of the species-level genome bins (SGBs) classification of Pasolli et al. ^28^, as they represent a greatly expanded set of genomes with thousands of new bacterial genomes that increase the number of mapped reads and allow better exploitation of metagenomic samples. We restricted ourselves to 3,127 SGBs that provide a good representation of human microbiome species diversity (**Methods**). We developed a method, Unique Relative Abundances (URA), for estimating the relative abundance of each SGB in every sample (**Methods**), which can be applied to any set of species, and provides better predictive power than Metaphlan (**Figure S1a-b, Methods**). Our method is based on examining only reads that map uniquely to a single SGB, since when using unique mappings we expect uniform coverage across SGB genome bins that have the same number of unique positions. This property allows robust estimation of relative abundances, as coverage across the genome bins depends linearly on the SGB’s relative abundance. The mean relative abundances of the different species are the same in the two Israeli cohorts but are somewhat different than in the US cohort (**Figure 1f-g, Table S1**).

### Microbiome diversity increases with age and associates with metabolic parameters

We first examined the association of microbiome diversity and human phenotypes since the literature is conflicted even on these basic associations. To this end, we computed alpha diversity using the species level Shannon index and ranked individuals by deciles of alpha diversity (**Figure 2a**). When comparing the top decile and the bottom decile of alpha diversity, we found that HbA1C%, BMI, fasting glucose and fasting triglycerides are significantly higher in the bottom decile while age and HDL cholesterol are significantly lower in the bottom decile (P-value < 10^−16^ after FDR correction, Mann Whitney rank-sum test), including a trend across deciles (**Table S2-S7**). Similarly, examining alpha diversity as a function of these traits, we found significant correlations between alpha diversity and each of these traits (**Figure 2b**). Notably, these associations were consistent in both the Israeli and U.S. cohorts (**Figure 2b**), even though the Israeli cohort has significantly higher alpha diversity values (**Figure 1e, 2b**, mean 7.3±0.77 vs. 7.18±0.67, P-value < 10^−40^,Mann Whitney rank-sum test). The higher diversity of the Israeli cohort persisted even when subsampling the Isareli cohort to match the U.S. cohort on age, gender and BMI (**Methods, Table S1**).

**Figure 2:**
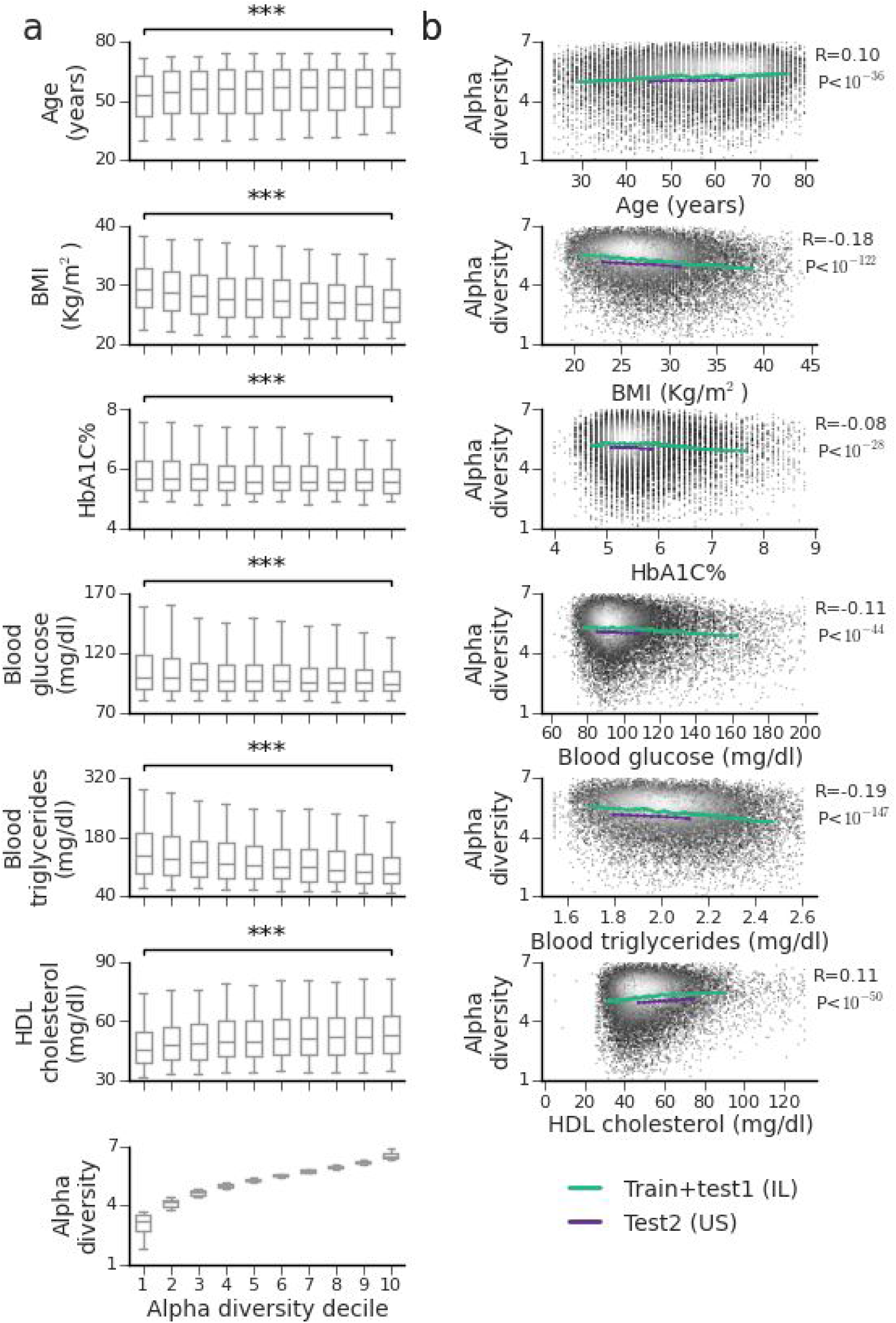
Species level Shannon alpha diversity significantly associates with many phenotypes. (a) A box-plot of the distribution of phenotype values, for each of 10 deciles of Shannon alpha diversity. Phenotype values in the first and last deciles of alpha diversity are compared using Mann-Whitney rank-sum test where *** signifies P value< 10^−16^ after FDR correction. Boxes correspond to 25-75 percentile of the distribution and whiskers bound percentiles 5-95. (b) Running average of alpha-diversity (y-axis) for the combined training and test1 cohort (green curve), and for the separate test2 cohort (purple curve), ordered by the phenotype values. For the larger Israeli cohort the average is on 1000 individuals with shift of 100 individuals; for the smaller U.S. cohort the group size and shift were chosen to obtain 10 points and the shift was 10% of the group size. The Pearson correlation and P-value shown are of the Israeli cohort, and are calculated on individual level data. Individual level data points are presented with grayscale colors. Data points with light colors have higher frequency.

### Microbiome-phenotype associations are consistent across continents

We previously employed linear mixed models to estimate the fraction of phenotypic variance that can be inferred from microbiome composition, termed *microbiome-association-index* (*b*^*2*^) ^*30*^. Our previous estimates were based on a cohort of 715 individuals and therefore had wide 95% confidence intervals, we revisited these estimates for our two new and larger cohorts. We estimated explained-variance based on alpha-diversity alone (**Figure 3a, Methods**), and based on the full species relative abundances (**Figure 3b, Methods**). We find that alpha diversity shows significant correlations, yet overall small explained variance, to different traits and we believe that literature inconsistencies between phenotype-alpha-diversity associations are a result of lack of power detecting these associations. Notably, our new estimates agreed well with the findings in our previous study (**Table S8**), but the current much larger cohort of 30,083 Israeli individuals gives substantially narrower 95% confidence intervals (**Table S9**). We found that microbiome composition strongly associates with self-reported diabetes (*b*^*2*^=52%), age (*b*^*2*^=28%), HbA1C% (*b*^*2*^=15%), fasting blood glucose (*b*^*2*^=13%), BMI (*b*^*2*^=11%), fasting triglyceride (*b*^*2*^=9%), HDL cholesterol (*b*^*2*^=6%) and smoking status (*b*^*2*^=6%). In contrast, the blood levels of thyroid-stimulating hormone (TSH), albumin and clotting (as measured by International Normalized Ratio, INR) were not significantly associated with the microbiome in our cohort. Notably, *b*^*2*^ estimates from our U.S. cohort of 3,974 individuals were consistent with those derived from the Israeli cohort (Pearson correlation R=0.75, P-value < 0.001) but had wider confidence intervals (**Figure 3b, Table S9,S10**). As expected, the variance explained by the full relative abundance matrix (our *b*^*2*^ estimates) was higher than that explained by alpha diversity alone, and furthermore these two microbiome features highly agreed (Pearson correlation R=0.52, P-value < 0.03).

**Figure 3:**
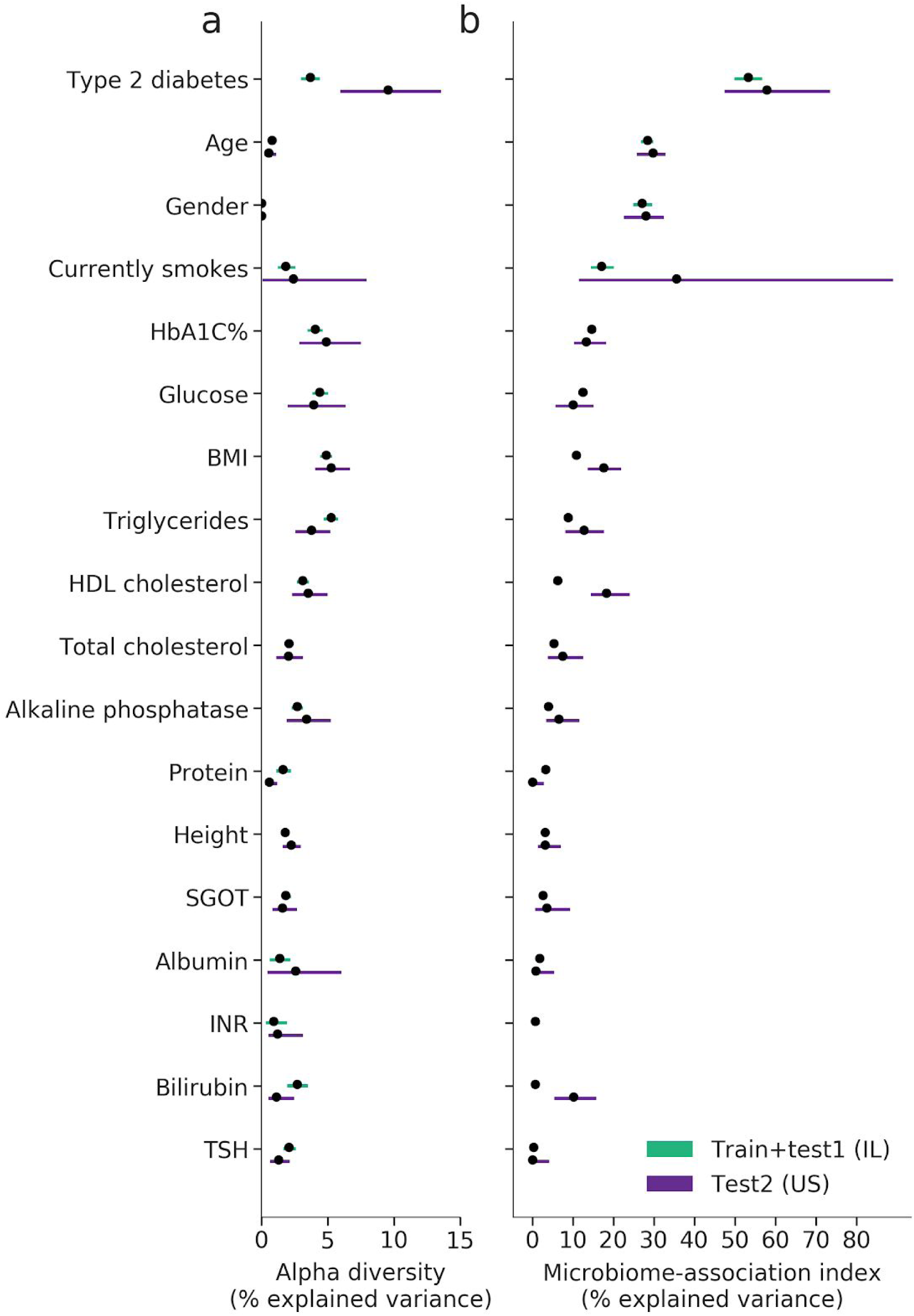
Explained variance of phenotypes based on microbiome features. (a) The proportion of variance of various phenotypes that can be explained using Shannon alpha diversity in the Israeli (green) and U.S. cohorts (purple) based on a linear model with covariates for age and gender. Also shown is the 95% confidence interval. (b) The proportion of variance of various phenotypes that can be explained using species-level relative abundances in the Israeli (green) and U.S. (purple) microbiome composition based on a linear mixed model estimation with covariates for age and gender (microbiome association index ^30^). Also shown is the 95% confidence interval. Estimates from the larger cohort have smaller confidence intervals.

### Different traits are accurately predicted by microbiome composition

We next asked whether various traits can be accurately predicted based only on microbiome composition. We compared two models; a linear model (with ridge regression regularization) and gradient boosted decision trees (GBDT) (**Methods**). Both models used only species relative abundances as input features to the model. Our models obtained significant predictions for many traits (**Figure 4a,b, Figures S2-3**) such as age (R^2^=0.21 for linear regression, R^2^=0.31 for GBDT, for 10 fold cross validation on train IL samples), gender (AUC=0.64, 0.78), HbA1C% (R^2^=0.15, 0.25) and BMI (R^2^=0.12, 0.15). Notebly, skin microbiome was recently shown to best predict age ^31^, and while our predictor based on gut microbiome is significantly more accurate than the reported gut microbiome predictor (Here R^2^=0.31 versus R^2^=0.17 by Huang et al. ^31^ skin microbiome predictor is far more accurate (R^2^=0.74 by Huang et al. ^31^).

**Figure 4:**
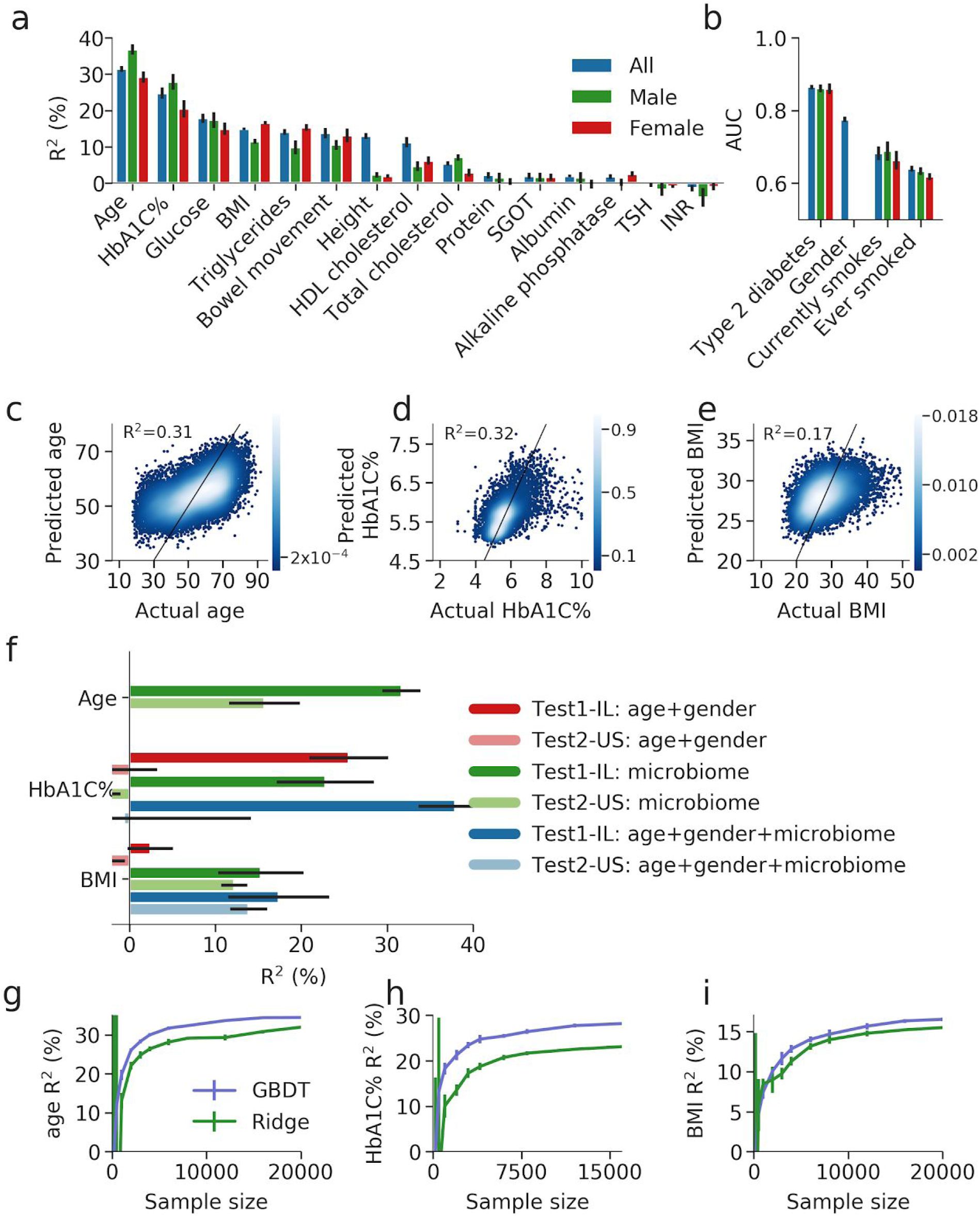
Prediction of phenotypes by the microbiome. (a) Coefficient of determination (R^2^) of GBDT prediction of different phenotypes based only on species level gut microbiome abundance. Results are obtained in a 10-fold cross validation scheme on the training set. Predictions are shown for three models, a model using the whole cohort, and a model for each gender. (b) Same as (a), but shown is the area under the curve (AUC) for predicting binary phenotypes. (c) - (e) Scatter plot of the phenotype and 10-fold cross-validation predicted values of the phenotype, for age, HbA1C% and BMI when training on the Israeli train cohort using GBDT. R^2^ of prediction is reported. Black line represents x=y. (f) Coefficient of determination (R^2^) of predictions of age, HbA1C% and BMI, for models trained with different sets of input features using GBDT, and tested on both the held-out Israel and U.S. test sets. Error bars of the test set are from bootstrapping. (g) - (i) Coefficient of determination (R^2^) and standard deviation error bars of predictions of age, HbA1C% and BMI obtained using GBDT (purple) or Ridge regression (green) models trained on sub-samples of the cohort train IL, of different sizes, and tested of the test IL cohort. For each cohort size *k*, 10 random sub-samples of *k* individuals were obtained and the mean and standard deviation of their predictions are shown.

We obtained significant predictions even when performing the analyses separately for each gender, with the exception of height which was significantly predicted (R^2^=0.13 GBDT) in the entire cohort but not in the gender-separated predictions, indicating that its predictions were driven by the prediction of gender. Since metformin, the most common drug used to treat patients with type2 diabetes, is known to affect microbiome composition, we also evaluated the performance of an HbA1C% predictor only on participants who did not report taking metformin and obtained equivalent performance (R^2^=0.19 GBDT).

The linear models are attractive since their accuracy was almost similar to boosting decision trees and they are easier to interpret. However, boosting trees performed better across 11 of 12 phenotypes (age being the exception) that had significant predictions (overall mean R^2^ improvement of 0.02+/-0.011, **Figure S1c-d**) suggesting that non-additive interactions between different bacteria are predictive of several traits. As additional evidence for the importance of non-additive interactions among bacteria in predicting traits, for both HbA1C% and BMI the R^2^ of the GBDT predictions on held-out subjects was higher than the estimated *b*^*2*^ for these traits (**Figure 3b**). As the *b*^*2*^ estimation used linear mixed models to estimate the fraction of variation predicted by the microbiome, it does not include any non-linear interaction captured by GBDT.

We investigated if the predictive power of the microbiome is mediated through age and gender, since some of the above traits such as HbA1C% are known to increase with age ^32^. We found that the microbiome composition predicted age with high accuracy (R^2^=0.32), and age and gender alone predict HbA1C% with R^2^=0.26 and BMI with R^2^=0.02 (GBDT, **Figure 4f**). Therefore, we asked whether microbiome-based predictions of HbA1C% and BMI are mediated entirely by its ability to predict age and gender, or whether it carried additional predictive power specific to these traits. We found that adding microbiome to age and gender to the GBDT model significantly improved the predictions of both HbA1C% (from R^2^=0.26 to 0.38, **Figure 4f**) and BMI (from R^2^=0.02 to 0.17, **Figure 4f**), demonstrating that some of the association between microbiome and these traits is not mediated through age and gender.

We also evaluated the accuracy of our above models, derived only based on the Israeli training set, on our two independent and held out cohorts from Israel and the U.S.. We found that microbiome base predictions for all traits are replicated in the Israeli held out cohort, and all microbiome base predictions except for that for HbA1C% are replicated in the U.S. cohort (**Figure 4f**), thereby validating the robustness of our models. We note that in general prediction accuracy was lower in the U.S cohort, which may be explained by differences in the microbiome composition between the IL and U.S. cohorts and by lower age and HbA1C% levels in the U.S. cohort (**Table 1**).

Finally, to examine the importance of cohort size on prediction accuracy, we applied our above prediction pipeline to different random subsamples of our training cohort, ranging from a few hundreds of subjects to 24,000 (**Methods**). We found that prediction accuracy increases with cohort size (**Figure 4g-i, S4**) and does not saturate even with a cohort of 1,000 individuals. For age, we observed an almost two-fold increase in the R^2^ (from 0.18 to 0.30) when increasing the cohort from 1,000 to 12,000 individuals. For cohorts of hundreds of individuals the standard deviation of the predictions was high, as in the case of HbA1C% for which different subsamples of 200 individuals can reach both R^2^=0.4 and R^2^=0.0 as likely outcomes (within 2 standard deviations). Together, these results highlight the need for obtaining large cohorts for microbiome studies, as is known to be the case in the field of human genetics.

### An atlas of bacterial species that robustly correlate with age, HbA1C% and BMI

We next sought to identify which individual bacterial species are responsible for driving the predictions of our models for age, HbA1C% and BMI, since these traits were predicted with the highest accuracy. We found many bacterial species that exhibited highly significant correlations to these traits (**Figure 5a-c, S5**, 967, 720, 1,056 bacteria out of the top 1,345 occuring bacteria had significant Spearman correlation, P-value < 0.05 after FDR correction for age, HbA1C% and BMI respectively, **Table S11-S16**). Moreover, the Spearman correlation of the bacterial abundances with these traits was in good agreement between the Israeli and U.S. cohorts (**Figure 5a-c, S5**, R=0.58, 0.50, 0.74 for age, HbA1C% and BMI, P-value<10^−87^). Notably, the 3 bacteria most strongly associated with BMI in both cohorts included two bacterial species from the *Eubacteriaceae* and *Clostridiaceae* family that was only recently assembled and that has no genome in public repositories (unknown SGB, **Figure 5a-c**).

**Figure 5:**
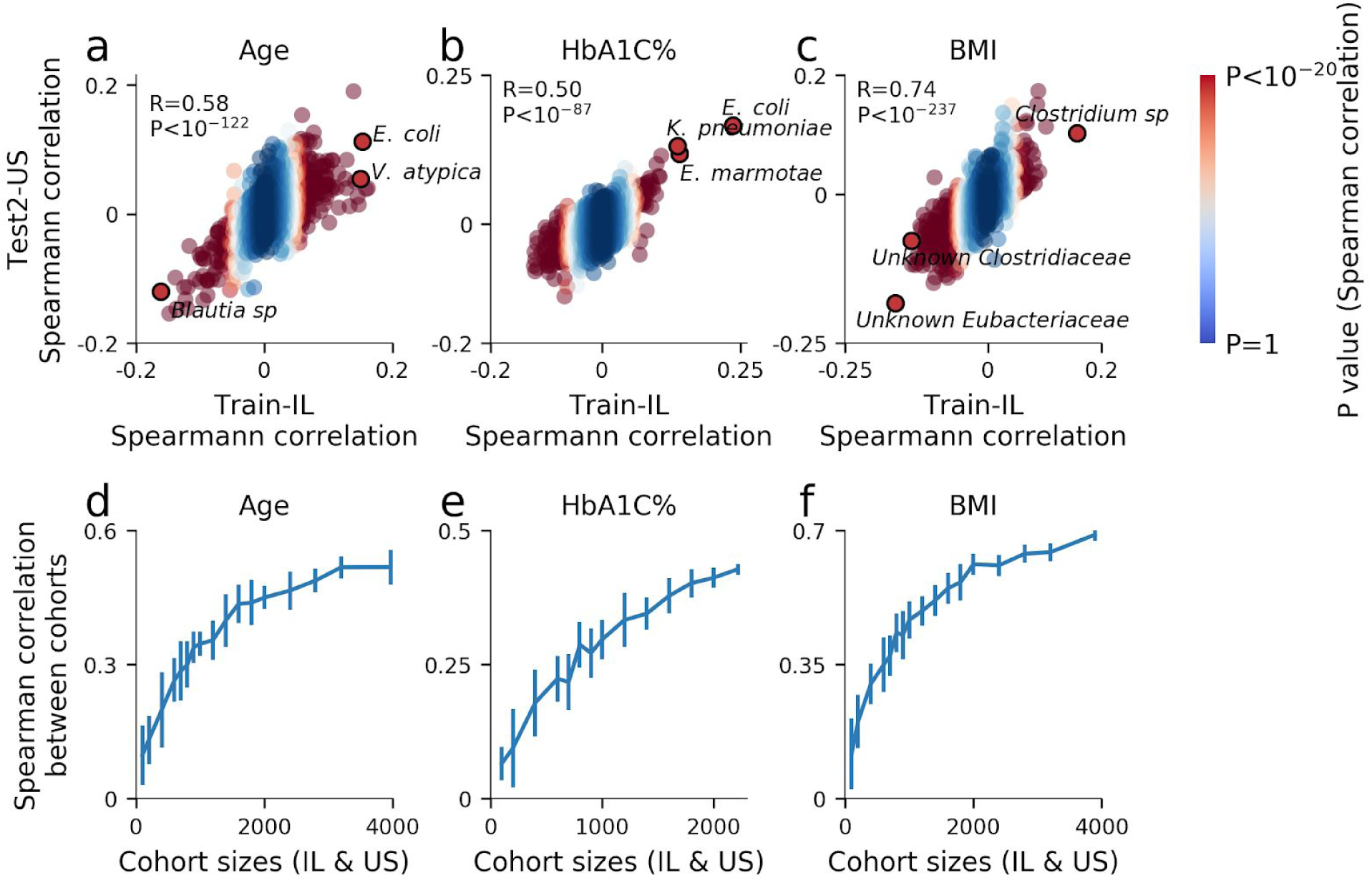
Predictive power of single species. (a) Spearman correlation of each bacterial species with age in the Israeli training cohort (x-axis, N=27,075) and the U.S. test cohort (y-axis, N=3,974). The correlation and P-value between the correlations of each cohort are shown. Bacteria are colored according to the P-values of the Spearman correlation in the Israeli cohort. The top three bacteria by Israeli P-values that replicate in the U.S. cohort are highlighted. (b) - (c) Same as (a) for HbA1C% and BMI. (d) Spearman correlation between the correlations of the Israeli cohort and U.S. cohorts as in (a) but for different sub-samples of cohort sizes. For each cohort size *k*, a sub-sample of *k* individuals was obtained from both the Israeli and U.S. cohorts and this procedure was repeated 10 times to obtain standard deviation error bars. (e) - (f) Same as (d) for HbA1C% and BMI.

Again, we subsample the cohorts to smaller cohort sizes and observe that large cohorts are necessary in order for results to replicate (**Figure 5d-f, Methods**).

### Functional characterization of gut microbiome

In order to identify bacterial mechanisms that can modulate metabolic phenotypes, we looked at module and pathway enrichments associated with collected phenotypes based on the URA microbiome composition (**Methods**). We found several modules and pathways which were significantly associated with human phenotypes (**Figure 6d-e, Table S17-S24**).

**Figure 6:**
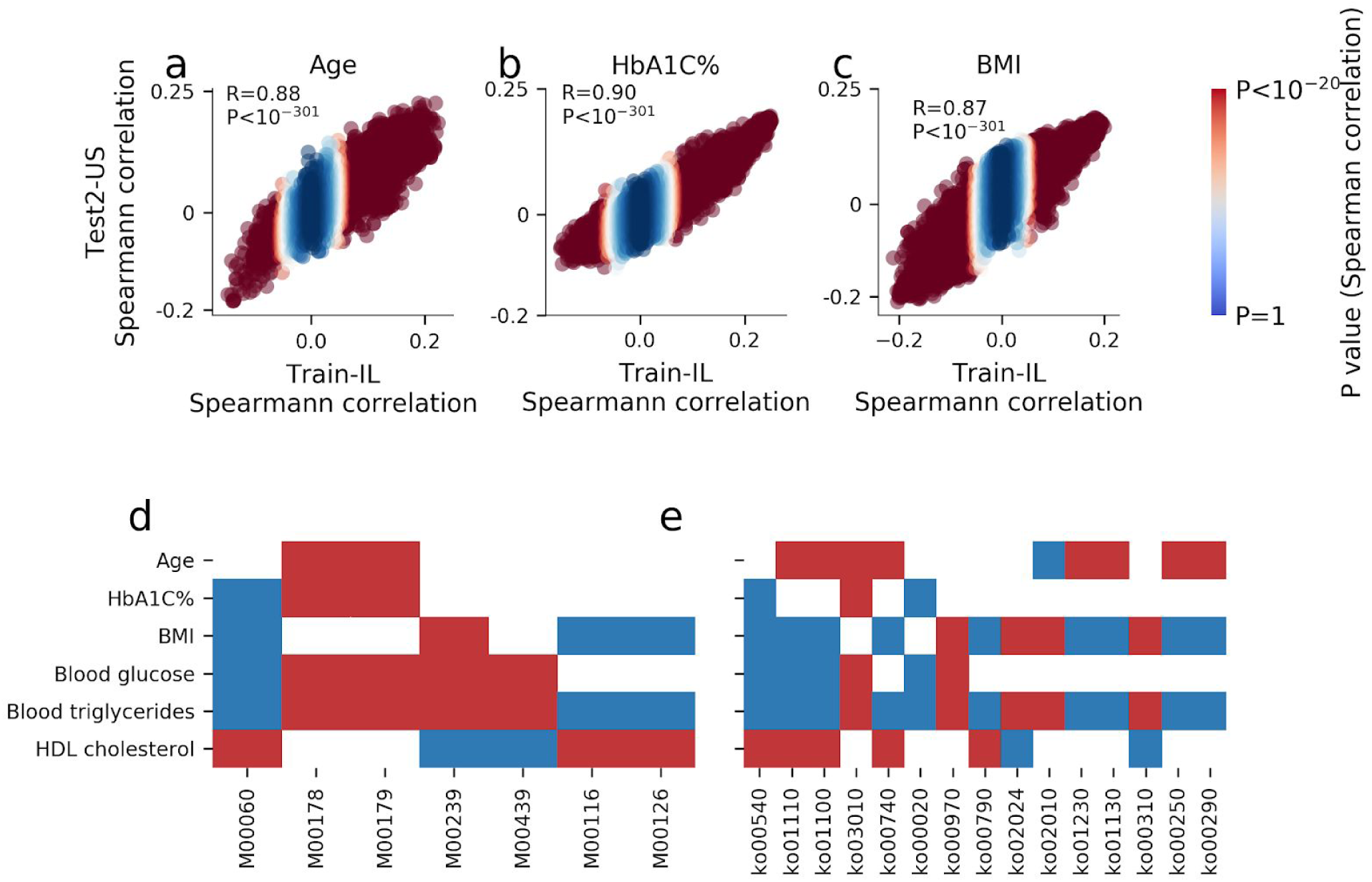
Functional analysis of modules and pathways. (a) - (c) Spearman correlation of each KEGG KO gene with age, HbA1C% and BMI in the Israeli training cohort (x-axis, N=27,075) and the U.S. test cohort (y-axis, N=3,974). The correlation and P-value between the correlations of each cohort are shown. Uniref are colored according to the P-values of the Spearman correlation in the Israeli cohort. (d) - (e) Heat map displaying only significant positive (blue) or negative (red) association between 6 phenotypes and KEGG KO modules (d) or pathways (e).

Most notably, the Lipopolysaccharide (LPS) biosynthesis pathway (ko00540) and the KDO2-lipid A biosynthesis module (M00060) were associated with adverse metabolic status (**Figure 6d-e**). Specifically, ko00540 and M00060 associated with increased levels of HbA1C% (P-values < 10^−3^ and 0.017 after FDR correction, respectively), BMI (P-values < 10^−3^ and 0.0014), fasting glucose (P-values < 10^−4^ and < 10^−2^) and fasting triglycerides (P-values < 10^−4^ and < 10^−3^) and with decreased levels of HDL-cholesterol (P-values < 10^−3^ and < 10^−2^). The LPS is a major component of the outer membrane of Gram-negative bacteria which is involved with toxicity, pathogenicity, antimicrobial resistance and other activities. LPS endotoxin purified from *E*.*coli* was previously shown to induce obese and insulin-resistant phenotypes when subcutaneously infused into mice ^33^. KDO2-lipid, the corresponding module, was shown to trigger defense-related responses by the human immune system to stimulate secretion of proinflammatory cytokines ^34^.

Another biological pathway that was positively associated with HbA1C% (P-value = 0.04), fasting glucose (P-value = 0.039) & fasting triglycerides (P-value = 0.034) is the citrate cycle (ko00020), which is an energy production metabolic pathway found mainly in aerobic bacteria. In concordance with this pathway, we found two modules that are part of the citrate cycle pathway (M00009 & M00011) that also positively associated with HbA1C% (P-value = 0.017 for both modules) and fasting glucose (P-value = 0.042 and = 0.044). The finding that this metabolic pathway is in significant association with pro-disease markers may be related to an oxygenic environment in the gut. This is in concordance with previous studies showing that chronic inflammation with oxidative gut-environment causes an imbalance between obligate and facultative anaerobes and supports gut dysbiosis ^35–37^.

We also found positive associations of vitamin metabolism modules, such as Menaquinone biosynthesis & Tetrahydrofolate biosynthesis (M00116 & M00126), with BMI (P-value = 0.019 and = 0.032, respectively), fasting triglycereides (P-value = 0.039 and = 0.04, respectively) and negative association with HDL-cholesterol (P-value = 0.026 and = 0.039), supporting previous findings of high concentration of menaquinone detected in the adipose tissues of obese adults ^38^. Menaquinone produced by *Enterobacter cloacae* had been shown to have strong correlation to BMI ^39^ and moreover, *E. cloacae* isolated from the gut of morbidly obese individuals induced obesity in germfree mice ^40^. Our analysis strengthens these links between bacteria producing Menaquinone and morbid obesity.

Finally, as a useful resource for the community, we compiled results into an atlas of summary statistics for all bacterial species and KEGG genes to 6 top predicted phenotypes (**Table S11-S22**). For each species and KEGG gene, we report bacterial associations to human phenotypes based on bacterial log relative abundances (**Methods)**. Specifically, we report the Spearman correlation coefficient and P-value, the Pearson correlation coefficient and P-value. For bacterial species we also provide the coefficient in the linear model (trained with Ridge regularization), and bacterial feature importance in the GBDT model using the feature attribution framework of SHapley Additive exPlanations ^41^ (SHAP). In genetics, summary statistics of single nucleotide polymorphisms are widely used to generate polygenic risk scores which were shown to be predictive of disease ^42,43^. Similarly, researchers can now use our resource to generate microbiome-based predictions of phenotypes in their datasets by extracting our reported bacterial regression coefficients and multiplying them by the log of the relative abundances of the corresponding species in their dataset.

## Discussion

In this study, we collected the largest cohort to date of metagenomic samples and phenotypic data from two continents, and analyzed it using a much expanded set of reference microbial species. Together, this allowed us to identify highly robust associations between gut microbiome composition and phenotypes, which replicate in both cohorts. We compiled these robust associations into an atlas that can be used by the community to derive trait predictions on smaller datasets, akin to the use of summary statistics in the field of genetics. We show that a large fraction of the variance of several traits such as age, HbA1C% and BMI can be accurately predicted by both linear models and boosting decision trees models. These predictions replicate across continents and there is also high agreement in the set of individual bacterial species that associate with these traits in the Israeli and U.S. cohorts.

When sub-sampling our large cohort into smaller sized cohorts, we found that even cohorts of 1,000 individuals have significantly lower average accuracy of associations between bacteria and phenotypes. Models derived from different sub-samples of smaller cohort sizes display high variability both in the set of bacteria that associate with each trait and in prediction accuracy. These results may explain the relatively low agreement that exists across studies in the set of bacteria associated with different traits and conditions, and they call for employing larger cohort sizes in microbiome studies.

Using an expanded reference set allowed us to study many bacterial species for the first time and to identify novel associations for them. Notably, even among the top associated bacteria we found unnamed bacteria that are prevalent and appear in hundreds and sometimes thousands of individuals from our cohort. This also provides a glimpse into possible functional drivers of some of these associations, which supports the hypothesis that metabolic syndrome phenotypes are strongly associated with low-grade subclinical inflammation leading to hyperglycemia and favoring the onset of type 2 diabetes and obesity ^44^. These findings emphasize the importance of expanding the reference set of the human microbiome even further, and suggest that such newly identified species may have strong associations with important host phenotypes.

Overall, by combining larger microbiome cohorts and expanded bacterial genome references we robustly characterize bacterial links to many important health parameters, serving an important first step towards unraveling the causal links and mechanisms by which bacteria affect host phenotype.

## Methods

### Recruitment

All participants are paying customers of a consumer microbiome company, who enrolled to get personalized algorithm based dietary recommendations, from January 2017 until January 2020. Exclusion criteria includes customers using antibiotics or antifungals three months prior stool sample collection, age under eighteen, pregnancy or less than three months post delivery, active fertility treatments or customers treated with short-acting insulin. We further excluded 2050 participants who self reported pancreatic disease, undetermined colitis, ulcerative colitis, crohn’s disease, inflammatory bowel disease, gestational diabetes or type1 diabetes. All participants provided a stool sample, and filled a medical questionnaire, before getting their dietary recommendations.

### Microbiome sample collection, processing and analysis

Participants provided a stool sample using an OMNIgene-Gut stool collection kit (DNA Genotek), and processed according to the methods described in Mendes-Soares et al. ^45^: Genomic DNA was purified using PowerMag Soil DNA isolation kit (MoBio) optimized for Tecan automated platform. Illumina compatible libraries were prepared as described in ^46^, and sequenced on an Illumina Nextera 500 (75bp, single end), or on a NovaSeq 6000 (100bps, single end). Reads were processed with Trimmomatic ^47^, to remove reads containing Illumina adapters, filter low quality reads and trim low quality regions; version 0.32 (parameters used: -phred33 ILLUMINACLIP:<adapter file>:2:30:10 SLIDINGWINDOW:6:20 CROP:100 MINLEN:90 for 100bps reads, CROP:75 MINLEN:65 for 75bps reads). Reads mapping to host DNA were detected by mapping with bowtie2 ^48,49^ (with default parameters and an index created from hg19) and removed from downstream analysis.

We used bowtie2 ^49^ to map samples from our cohort versus an index built from the set of representatives of the SGBs (demanding all mappings of length 100/75 to score -40 or above). On average 77 percent of reads (minimum of 50 percent, maximum of 86 percent) mapped to bacterial representatives. This mapping percentage is in line with the original mappability estimates of Pasolli et al. ^25^ for westernized gut microbiome samples on the set of representatives of the reference set.

All samples of 75bps (Nextera 500) were subsequently downsampled to a depth of 8M mapped reads, and all samples of 100bps (NovaSeq 6000) were subsequently downsampled to a depth of 5M mapped reads. That is, sub-sampling was performed after mapping to a bowtie2 index of the bacterial reference dataset, and only reads which mapped to one or more bacteria in the dataset were taken. For samples with fewer mapped reads, all mapped reads were taken, this accounted for about half of the 75bps samples, and less than 5% of the 100bps samples.

All atlas correlations and all predictions were performed also on the subset of samples of 75bps reads with 8M mapped reads or of 100 bps reads with 5M mapped reads. All results replicate, but with lower power, since they were performed on less samples.

### Relative abundance estimation of SGBs - Unique Relative Abundance (URA)

The bacterial reference dataset for relative abundance estimation is based on the representative assembly of the species-level genome bins (SGBs) and genus-level genome bins (GGBs) defined by Pasolli et al. ^28^. By construction, all assemblies in each SGB are at high average nucleotide identity with one another, and the representative was chosen to be the best quality assembly amongst them.

Out of the 4,930 human SGBs (associated with various body sites), we chose to work with 3,127 SGBs, which were characterized by either belonging to a unique genus or with at least 5 assemblies to justify having a new SGB. We employed this restriction, since we noticed that the cutoff threshold used by Pasolli et. al. to cluster assemblies into SGBs resulted in small groups with little nucleotide difference from a large nearby SGB is artificially split to a new SGB.

Abundance was calculated by counting reads that best matched to a single SGB of the set. In order to avoid sample reads which may be assigned to more than one SGBs (which might mislead us to believe an SGB appears in a sample when it actually does not), we created a mapping of all 100/75-bps reads which are unique to a single of these representatives. We divided each representative genome assembly to consecutive windows such that each window includes 100 unique 100/75-bp reads (unique-100-bins). Since different proportions of reads are unique in different areas of the assembly, these windows are not of constant length, but the number of sample reads expected to uniquely map to them should be constant.

We used bowtie2 ^49^ to map samples from our cohort versus an index built from the set of representatives of the SGBs (demanding all mappings of length 100/75 to score -40 or above). When analysing the mapping, we looked only at reads whose best map is unique (thus mapped to a location which is unique in the set of representatives). We count the number of reads uniquely mapped to each window of each SGB.

To assess the cover of each SGB, we first choose a window size, which is a multiple of the original unique-100-bins, for which the average cover is at least 20 reads. Next, we sum the number of reads in this enlarged-window, and test the distribution of covers over the windows.

Finally, we take the dense mean of that distribution ^50^, in order to avoid our cover estimation being biased by a relatively small part of the reference which is highly covered (may come about from plasmids or horizontal transfer which was not identified in the uniqueness process since it did not appear in any other representative) or lowly covered (since this is a representative of an SGB, a strain present in our sample may not include all parts of the representative). When the dense 50% of the cover distribution starts above 0 we conclude the SGB exists in the sample, and we estimate its relative abundance. The cover estimation for each SGB is the dense mean cover of its representative, normalized by the enlarged-window size.

The relative abundance estimation is the cover divided by the sum of the covers of all representatives we concluded exist in this sample.

Code of the algorithm, and for building the necessary databases, is provided in github: https://github.com/erans99/UniqueRelativeAbundance

### Data and code availability

All of the source code used to generate the analyses of this paper will be made available upon publication, including a subset of 100 raw metagenomic samples on which the code can be run to validate the pipeline and reproduce the results obtained on these samples. In addition, as is commonly done in the field of human genetics, full tables of summary statistics of the correlations between bacterial species abundance and human traits will be made available. As in the field of human genetics, access to the raw metagenomic sequencing reads of all samples cannot be provided due to restrictions set forward in the informed consent

### Predictive Power of the Unique Relative Abundance (URA) Method

To evaluate our unique relative abundance method (URA), we tested the quality of predictions derived from MetaPhlan ^27^ species relative abundances, vs. our URA abundances.

In order to discriminate between the predictive power derived from the estimation method vs. that coming from the expanded set of species, we created a new URA database, including only the subset of 998 SGBs that Pasolli et al. marked as known NCBI human bacterial species. We find that URA with this reference set achieves a higher prediction accuracy than MetaPhlan ^27^ for different phenotypes, and for most phenotypes a lower power than the prediction accuracy using the expanded 3,127 SGBs reference set (**Figure S1a-b**).

### Strengths and limitations of Unique Relative Abundance (URA) method

The URA method is based on taking a set of reference genomes, one representing each of the different species, and looking only at the reads that are unique to a single one of these representatives. This mapping is weighted against pre-processed found uniqueness between bacterial species in the reference dataset (UREF). The expectation is that reads are randomly sampled from the gut bacterial population. In order to fulfill this assumption the implemented method uses only single end reads.

While this process gives a good estimation of abundances, as can be seen by its high predictive power (**Figure S1a-b** shows comparison with predictive power of Metaphlan abundances), it does not mean that any read mapped to a species representative is necessarily a read originating from that species. Specifically, deletions, copy number variations and horizontal gene transfers may cause some of the reads originating for one bacteria in some individual’s gut, to be assigned to another bacteria’s UREF. Thus, the method is appropriate for species abundance estimation, but requires careful interpretation for gene level analysis.

### Cohort matching

We subsampled the IL cohort to match the U.S. cohort on age, gender and BMI, using the MatchIt package from CRAN repository for r.

### Alpha diversity explained variance

We calculated the alpha diversity explained variance by regressing out gender and age from each phenotype, and then using ordinary least squares modeled the phenotype by alpha diversity. To get confidence intervals, we bootstrapped the data 10,000 times.

### Microbiome-association-index

We calculated *b*^*2*^ estimates using linear mixed models as was previously described ^30^. We used age and gender as fixed effects covariates, and built a microbiome genetic-relationship-matrix, using our developed SGB based relative abundances. The *b*^*2*^ calculation assumes that the phenotype distributed normally, we removed sample outliers from the IL and US cohorts using the same thresholds (removing less than 5% of individuals Table **S25**). To account for differences between the population and study prevalence of binary traits, we applied the correction of Lee et al. ^51^ which has been shown to provide a lower bound on the fraction of explained variance ^52^. We also provide uncorrected estimates in Table **S26**. Phenotype distributions of blood SGPT levels were far from normally distributed and were not estimated.

### Phenotypes prediction

We used the gradient boosting trees regressor from Xgboost ^53^ as the algorithm for the regression predictive model for different phenotypes. We used the gradient boosting trees classifier from Xgboost as the algorithm for the classification predictive model for phenotypes with binary values. All hyperparameters of the xgboost were fitted based only on cross validation of the train set.

The parameters of the predictors when using microbiome features were: colsample_bylevel=0.075, max_depth=6, learning_rate=0.0025, n_estimators=4000, subsample=0.6, min_child_weight=20. These parameters were used for regression as well as classification.

The rest of the parameters had the default values of Xgboost.

For the Ridge linear regression we used the RidgeCV from the scikit-learn package. The parameters used for the regressor were: alphas=[0.1,1,10,100,1000], normalize=True. The rest of the parameters were the default. The input to the Ridge linear regression was log transformed SGB abundance.

For binary phenotypes SGD classifier from the scikit-learn package was used, with default parameters (L2 normalization).

When using microbiome features for the prediction, only the top 1345 occuring SGBs were used, i.e., the SGBs that were found in at least 5% of the samples, to avoid overfitting on rare SGBs.

### Calculating prediction accuracy as a function of cohort size

For cohort size n (for n=200, 500, 1000, 2000, 3000, 4000, 6000, 8000, 12000, 16000, 20000, 24000; for prediction of HbA1c the maximum size was 16000) we repeated the following process 10 times: we randomly selected a subset of n samples, built a predictive model for the phenotype from the subset of samples, and tested its predictions on the independent test sets (test1-IL and test2-US test-sets described in main text), and calculated the accuracy (R square) of the prediction. By repeating the procedure 10 times we received the mean and standard deviation of the prediction accuracy estimate, for each IL trained subset size.

### Predictive power of single species

For cohort size n (for n=100, 200, 400, 600, 700, 800, 900, 1000, 1200, 1400, 1600, 1800, 2000, 2400, 2800, 3200) we repeated the following process 10 times: we randomly selected a subset of n samples from each cohort, calculated the Spearman correlations of each species with the phenotype in each cohort, and correlate the Spearman correlations of the two cohorts, over all species (not just those that pass a threshold p-value). We plot the mean and standard deviation of this correlation, as calculated over the 10 repeats.

### Correlation of phenotypes with biological annotation

Gene prediction, and annotations, of the representative genomes of the SGBs were performed in advance by Pasolli et al., including assigning Kegg Orthologies (KOs) to identified genes. To calculate the pseudo-abundances of each KO in each member of the cohort we multiplied the relative abundances of the SGBs with the number of times each KO appears in the representative genome.

For each KO which appears in at least 5 positions in the representatives, the vector of its pseudo-abundance (one per individual in our cohort) was correlated versus different phenotypes (an atlas is provided in **Table S17-S22**), and then ranked, from the most significantly negatively correlated to the most significantly positively correlated with the phenotype. This process was performed for each cohort separately, and shows significant concordance between the IL and the US cohorts (**Figure 6 a-c**).

After ranking the KOs against a phenotype, ranksum analysis was used in order to search for pathways and modules which are significantly enriched or significantly diminished (**Table S23-S24**), with connection with the phenotype.

## Supplementary Figures

**Figure S1:**
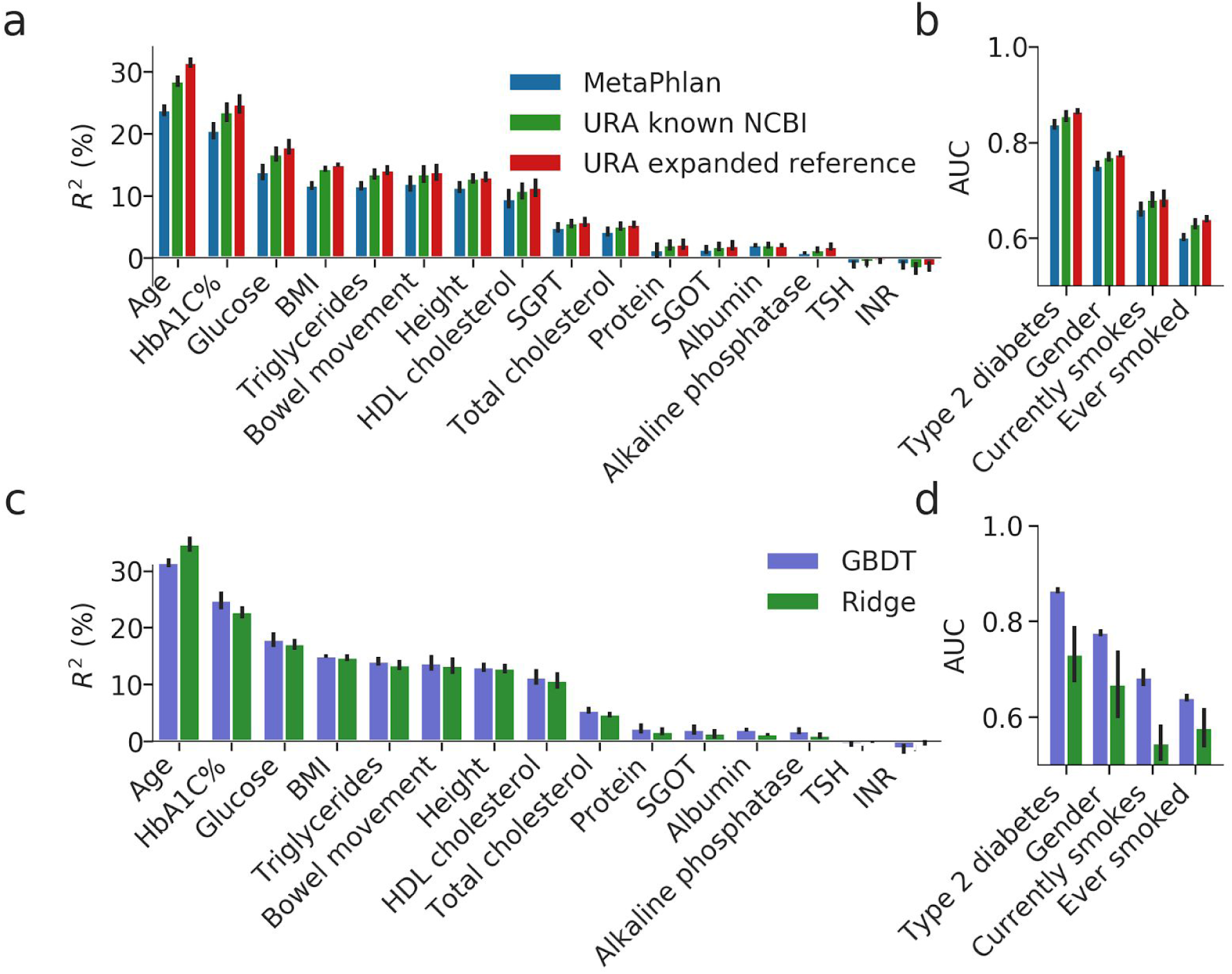
Comparisons of prediction of phenotypes by the microbiome. (a) Comparison of the predictive power of three different sets of species level abundance estimations of gut microbiome. In blue, predictions are performed using the baseline Metaphlan species abundances, in green predictions are performed on abundances calculated using the URA algorithm on the sub-set of SGBs that were known prior to the work of Pasolli et al. ^28^, and in red predictions are performed on URA abundances. For all three, the coefficient of determination (R^2^) of GBDT prediction of different phenotypes are obtained in a 10-fold cross validation scheme on the training set. (b) Same as (a), but shown is the area under the curve (AUC) for predicting binary phenotypes. (c) Comparison of the predictive power of GBDT model (in blue) versus Ridge regression model (in green). Coefficient of determination (R^2^) of prediction of different phenotypes based only on species level gut microbiome abundance. Results are obtained in a 10-fold cross validation scheme on the training set. (d) Same as (c), but shown is the area under the curve (AUC) for predicting binary phenotypes.

**Figure S2:**
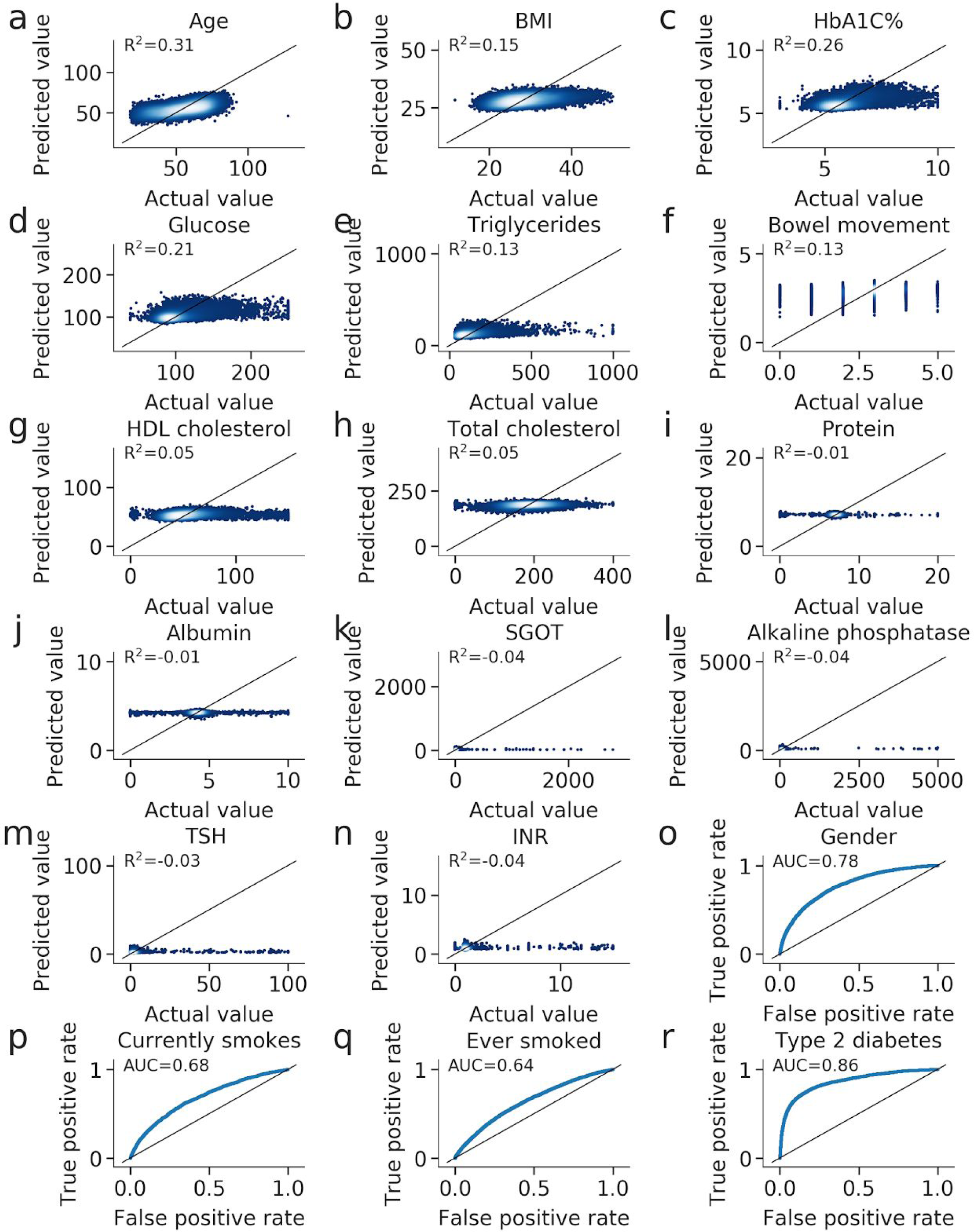
GDBT prediction of phenotypes by the microbiome. (a) - (n) Scatter plots of 10-fold cross-validation predicted values of quantitative phenotypes when training on the Israeli train cohort using GBDT. R2 of prediction is reported. Black line represents x=y. (o) - (r) ROC curve plots of 10-fold cross-validation predicted values of binary phenotypes when training on the Israeli train cohort using GBDT. AUC of prediction is reported. Black line represents x=y.

**Figure S3:**
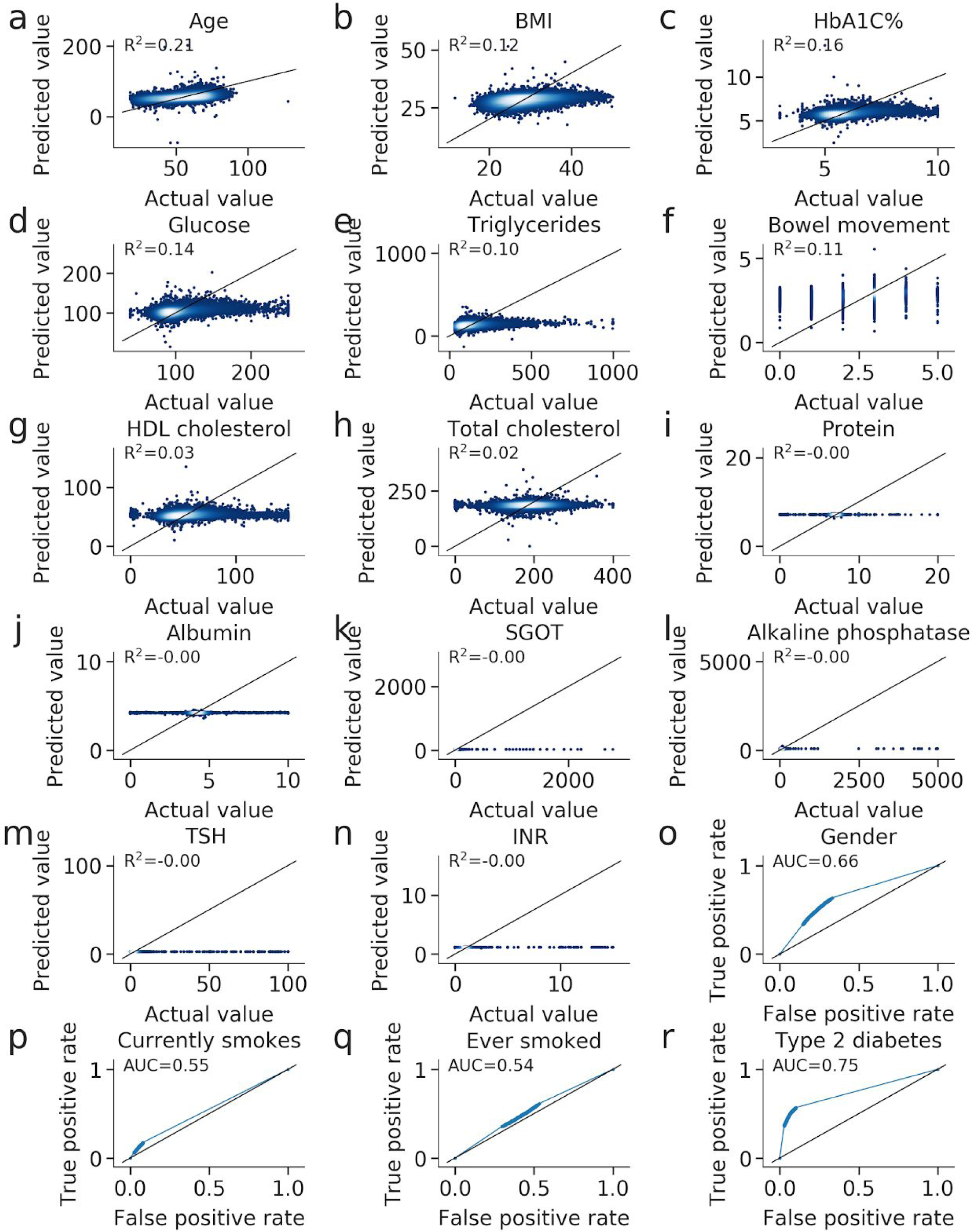
Prediction of phenotypes by the microbiome using Ridge regression.-. (a) - (n) Scatter plots of 10-fold cross-validation predicted values of quantitative phenotypes when training on the Israeli train cohort using Ridge regression. R^2^ of prediction is reported. Black line represents x=y. (o) - (r) ROC curve plots of 10-fold cross-validation predicted values of binary phenotypes when training on the Israeli train cohort using Ridge regression. AUC of prediction is reported. Black line represents x=y.

**Figure S4:**
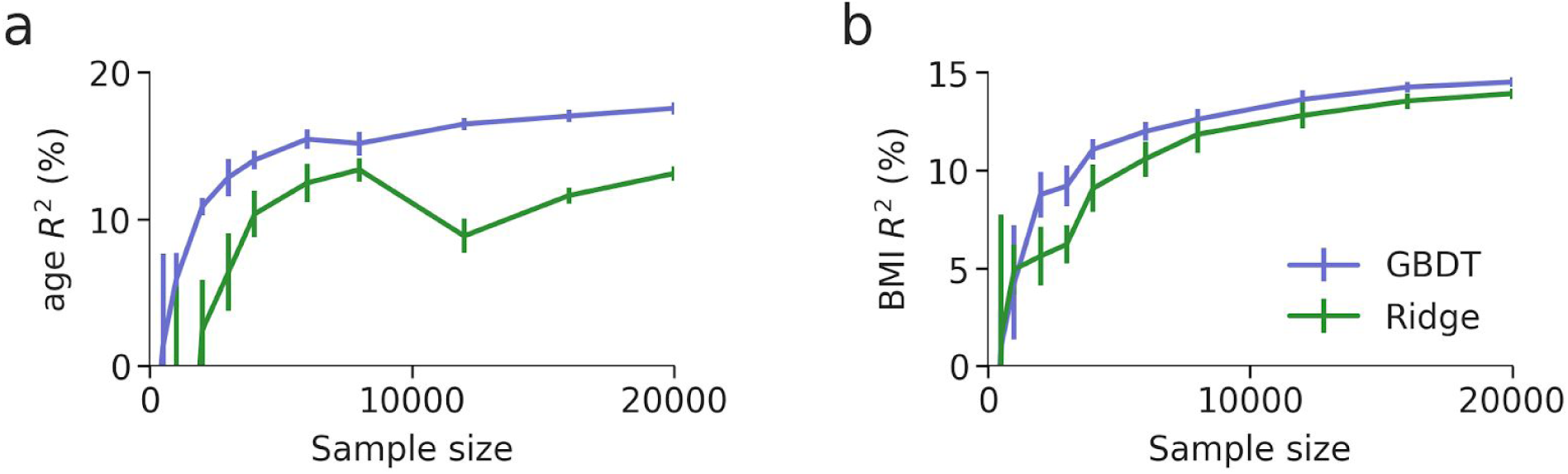
Prediction trained on sub-sampled IL cohort sizes using GBDT or Ridge tested on the US test cohort. (a) - (b) Coefficient of determination (R^2^) and standard deviation error bars of predictions of age (a) and BMI (b) obtained using a GBDT (purple) or Ridge regression (green) models trained on sub-samples of the cohort train IL, of different sizes, and tested of the whole test US cohort. For each cohort size *k*, 10 random sub-samples of *k* individuals were obtained and the mean and standard deviation of their predictions are shown.

